# Guidance of zoospores by potassium gradient sensing mediates aggregation

**DOI:** 10.1101/470864

**Authors:** Eric Galiana, Celine Cohen, Philippe Thomen, Catherine Etienne, Xavier Noblin

**Affiliations:** Université Côte d’Azur, INRA, CNRS, ISA, Sophia Antipolis, France; Université Côte d’Azur, CNRS UMR 7010, Institut de Physique de Nice, Parc Valrose, 06108 Nice, France

**Keywords:** *Phytophthora*, zoospore, Negative chemotaxis, Bioconvection, Aggregation

## Abstract

Biflagellate zoospores of some phytopathogenic *Phytophthora* species spontaneously aggregate within few minutes in suspension. Depending on species auto-aggregate formation results from bioconvection or from a sequence bioconvection-positive chemotaxis. In this study, we show that *P. parasitica* zoospores may form aggregates upon application of a K^+^ gradient in particular geometric arrangements. Based on the use of macro- and microfluidic devices, in addition to time-lapse live imaging both in the vertical and horizontal planes, we defined (i) the spatiotemporal and concentration scale evolution within the gradient in correlation with (ii) cell distribution and (iii) metrics of zoospore motion (velocity, trajectory). The results indicated that K^+^-induced aggregates result from a single bi-phasic temporal sequence involving negative chemotaxis and then bioconvection in a K+ gradient concentration scale [0-17 mM]. Each K^+^-sensing cell undergoes a forward-to-backward movement within a threshold range of 1-4mM, thereby forming progressively a swarm. Once a critical population density is achieved zoospores form a plume which undergoes a downward migration leading to aggregate formation on the support surface. We discuss putative sources for K^+^ gradient generation in natural environment (zoospore population, microbiota, plant roots, soil particles), and implication for events preceding inoculum formation on host plant.

## 1. Introduction

Within Oomycetes the *Phytophthora* genus includes some of the most destructive plant pathogens to manage causing destructive diseases on crops and natural ecosystems worldwide. They grow as filamentous coenocytic hyphae and move mainly as unicellular zoospores. Zoospore motility is critical for successful infection. To reach potential hosts a flagella-mediated random motility allows zoospores to explore environment much more efficiently than spreading by Brownian motion alone. The locomotor apparatus consists in two flagella, a whiplash one directed posteriorly and a tinsel one directed anteriorly [1, 2]. Flagella assembly and function appear to depend at least on a Gα-subunit protein and a bZip transcription factor and dynein-based molecular motors [3, 4].

A bias on the random motility may be imposed by exogenous signals. Zoospores possess sensory systems which confer the ability to respond to chemical gradients (chemotaxis), oxygen (aerotaxis), ionic fields (electrotaxis) or light (phototaxis). To target plant tissues they detect gradients of a variety of compounds including ions, plant isoflavones, amino acids, sugars [5-7]. For example, zoospores of *Phytophthora palmivora* exhibit anodal electrotaxis in electrical fields ≥0.5 V/m comparable in size to the physiological fields around roots [8]. When a zoospore reaches a potential infection site, the cell drops its flagella, synthesizes a primary cell wall and produces a germ tube before host tissue penetration [1, 9-11].

Zoospores also exhibit the ability to control their motility in response to self-produced signals. They constitute swarm or swim towards encysted spores leading to auto-aggregation (autotaxis) [12, 13] or to biofilm formation on plant surface [14, 15]. The incidence of such multicellular structures on plant infection has been largely neglected, the pathogenic process being considered until recently at the unicellular level for investigating the interaction with the host. Some data now support the hypothesis according to which *Phytophthora* ability for a multicellular behavior contributes to the dynamics of the interaction with a host through interspecific or intraspecific cell-to-cell signaling [14, 16]. How zoospores perceive cell density, transduce signal(s) and govern auto-aggregation is poorly understood. Mathematical and experimental data indicate that auto-aggregation in *Phytophthora* species results from bioconvection leading to plume formation [17] or from a sequential combination of bioconvection and then chemotaxis between plumes [13]. It is well described that *Phytophthora* species produce and use molecules to monitor zoospore density but no auto-attractant has been identified so far. The *P. parasitica* zoospores produce an AI-2-like signal (but not N-Acyl homoserine lactones) which could drive quorum sensing [18]. They also secrete cAMP a putative chemoeffector during biofilm formation [19].

Behavior of zoospores in response to change in ionic conditions suggests that cationic fluxes could be involved in collective motion but their nature and their role remain to be established. Ca^++^ plays a central role in autonomous encystment, adhesion, germination and auto-aggregation [12] but does not directly trigger cooperative behaviors of zoospores and acts more like a secondary messenger [18, 20]. K^+^ homeostasis influences the locomotion and encystment of zoospores. High external concentrations (5-10 mM) of potassium salts reduce the swimming speed of and cause the *P. palmivora* zoospores to swim in a jerky fashion [5]. Potassium sensing provokes negative chemotaxis of zoospores of *Phytophthora palmivora* [6].

Seeking to characterize which signals may alter zoospore random motility and initiate auto-aggregation or biofilm formation, we got evidence that aggregation of *P. parasitica* zoospores is elicited through perception of monocationic gradients in a particular geometric arrangement. Using macro- and microfluidic systems combined with time-lapse live imaging for measuring fluorescent intensity of impermeant cation probes we defined characteristics of K^+^ gradients and zoospore motion.

## 2. Material and methods

### 2.1 Zoospore suspension preparation

Mycelium of *Phytophthora parasitica* (isolate 310, *Phytophthora* INRA collection, Sophia-Antipolis) was routinely cultured on malt agar at 24 C in the dark. For zoospore production, mycelium was grown for one week in V8 liquid medium at 24°C under continuous light. The material was then drained, macerated and incubated for a further four days on water agar (2%). The zoospores were released by a heat shock treatment: incubation for 30 min at 4^⍰^C followed by incubation for 20 minutes at 37°C. Ten milliliters (per Petri dish Φ100mm) of 2mM Mes [2-(N-Morpholino) ethanesulfonic acid]-NaOH buffer pH 6,5 was added between incubations. Excepted when mentioned zoospore concentration was calibrated to 5.10^5^ cells/μl.

### 2.2. Droplet assay

Three chemotaxis devices were used for zoospore motion studies. A first one was the droplet assay. A controlled profile of the signal was generated by applying a [1mM-1M] gradient in a local and oriented-manner to a freshly prepared suspension of zoospores so as to reach a more or less steady profile by diffusion. The basic operation involved microinjection of putative chemo effectors (1/200, V/V) at the periphery of droplets (200-500 μL) including zoospores and deposited on glass sides or PDMS (Polydimethylsiloxane) stamps. The qualitative and quantitative zoospore response occurred by analyzing chemoeffector-induced cell trajectories. Various conditions (ionic composition and strength, pH, nutritive resource, cell density) were assayed leading to the identification of K^+^ and Na^+^ sensing as a primary signal for zoospore aggregation.

### 2.3. Passive dispersion system used to generate ionic gradient

A passive dispersion system was used to generate a diffusion gradient for tracking simultaneously extracellular concentration of potassium and zoospore distribution. Cells were preloaded with 2μM Asante Potassium Green-2 (APG-2 TMA+ salt, Teflabs, 3622), a potassium-specific and non-permeant fluorescent dye. Fifty microliters of cell suspension were placed in a μ-chamber (μ-Slide VI - Flat, Ibidi size l:17 mm; w: 3.8 mm; h: 400 μm) before application of 0.5 μl of 500mM KCl. At different time points, each μ-chamber content was observed using a confocal microscope (LSM 880-Zeiss) and the Tile Scan tool. Image sizes of 1.2 mm x 1.2 mm were generated from the application point and along the length. APG-2 was excited with a 488nm argon laser and the fluorescence emission was captured using the channel mode (band pass 510-590nm). The distant between the application point and the upper position in the chamber suitable for zoospore motion was determined using the transmission-photomultiplier. The first 1,2mmx0,4mm corresponding to this position was divided in 3 technical replicates (0,4×0,4) to measure mean APG-2 fluorescent intensity in each area. The values of higher ionic concentration suitable for zoospore motion was calculated from the difference between APG-2 FI measured at this position, after and before potassium application, and in reference to FI values measured with a range of discrete concentrations. Image analyses were done with the ZEN 2 software (Zeiss).

### 2.4. Microfluidic device

The third device dedicated to capture immediate response of zoospores at the single cell level consisted in a PDMS microfluidic circuit for submitting zoospores to a continuous flow presenting a gradient of potassium concentration. The device is presented on Figure 5A). Three channels (h*x w* = 0.05 mm x 0.1 mm) intersect as a cross and fuse into a last channel with much larger width (HxW=0,2 mm x 1 mm) which is called the chemotactic chamber where the spores will be observed. Spores are injected in the middle inlet and the 100 mM KCI solutions injected in the two lateral channels. The device was fabricated using soft lithography techniques [21] all the steps being realized in the clean room of Institut de Physique de Nice. Molds in SU-8 exposed at a resolution of 50800 DPI are covered with 10:1 Sylgard PDMS to be cured. After unmolding and puncturing for inlets, plasma bonding on clean glass slides is used to seal the channels. Teflon tubing directly inserted in the PDMS holes are used to connect the solutions reservoirs (2 ml) to the system. Liquid flows are driven using a pressure controller (Fluigent) in the range 0-100 mbar between the three inlets and the outlet which is at atmospheric pressure. The main interest of this controller is that changes of pressure can be done rapidly, allowing to reach quickly the desired conditions around the spores in the chamber. A high-speed camera (Phantom v7.11) was used and placed on an inverted microscope Zeiss Axiovert 200M with a x10 objective to acquire movies at a speed of 200 fps.

### 2.5. Microscopy for image acquisition along horizontal or vertical axes

Zoospore motion was captured using different instruments of the Microscopy Platform-Sophia Agrobiotech Institute (INRA 1355-UCA-CNRS 7254-INRA PACA Sophia Antipolis): the digital microscope VHX-2000 (Keyence); the inverted confocal microscope LSM 880 (Zeiss); the axioImagerZ1 (Zeiss) equipped for bright and epifluorescence, as well as with the AxioCam MRm camera. An Axioskop (Zeiss) microscope mounted vertically enables imaging of zoospores moving in a vertical plan. To achieve this, the microscope was rotated by 90°, appropriately supported on the back of its frame along the horizontal axis, and with its stage properly positioned with a polystyrene serving as a chock. Using these instruments, movie acquisition was controlled by 3D profile VHX-H3M (Keyence) or ZEN (Zeiss) software and resulted to sequences from 10 to 30 seconds at 20-30 frames per second.

### 2.6. Image analysis

Dynamics of zoospore motion was first investigated by single-particle tracking with different image-processing algorithms available as plugins in the ImageJ or Fiji software libraries. Prior further analysis, processing of phase contrast zoospore consisted to TIFF format conversion, image inversion, threshold adjustment and binary conversion. The Velocity_Measurement_Tool plugins enabled to generate kymographs. The MosaicSuite [22] allowed us to define trajectories. Image size was adjusted as cells corresponded to 3 to 5 pixel particles. The TrackMate plugin [23] was used to examine motion metrics. Frame sequences were analyzed by a time step of two seconds [Initial processing used criteria: estimated blob diameter, 10μm; Auto initial thresholding; linking max distance, 30 μm; gap-closing max distance, 15 μm; gap-closing max frame,1]. Utilizing generated track statistics, only paths defined continuously for at least 1,5s were selected for further treatment of TRACK_MEAN_SPEED data. Manual Tracking (https://imagej.nih.gov/ij/plugins/track/track.html) was used to analyze high cell density or the linear-rotational transition.

## 3. Results

### 3.1. Potassium and sodium induces zoospore aggregation in a concentration-specific manner

The droplet assay (Figure 1A) was first used both to evaluate specificity level and to determine optimal conditions for detection of ion gradients by zoospores. Upright microscopes were used for observation in the horizontal plane (Figure 1A). A microscope setup was adjusted in a way to position the axis of the lens horizontally for observing displacement in the vertical plane (Figure 1B). With appropriate ion concentration, three states can be defined and reproducibly observed (Figure2A): (*i*) a free state (FREE) corresponding to untreated cells which are uniformly distributed in the droplet and exhibited random motion; (*ii*) a swarm state (SWA), accounting for ion treated cells, during which the vast majority of zoospores migrated upward and gradually constituted a swarm with increasing cell density in a progressively restricted area within a few minutes; (*iii*) an aggregate state (AGG) corresponding to zoospores forming a plume when a critical population density is achieved, leading to a downward migration and then to cell gathering on the support surface.

**Figure 1:**
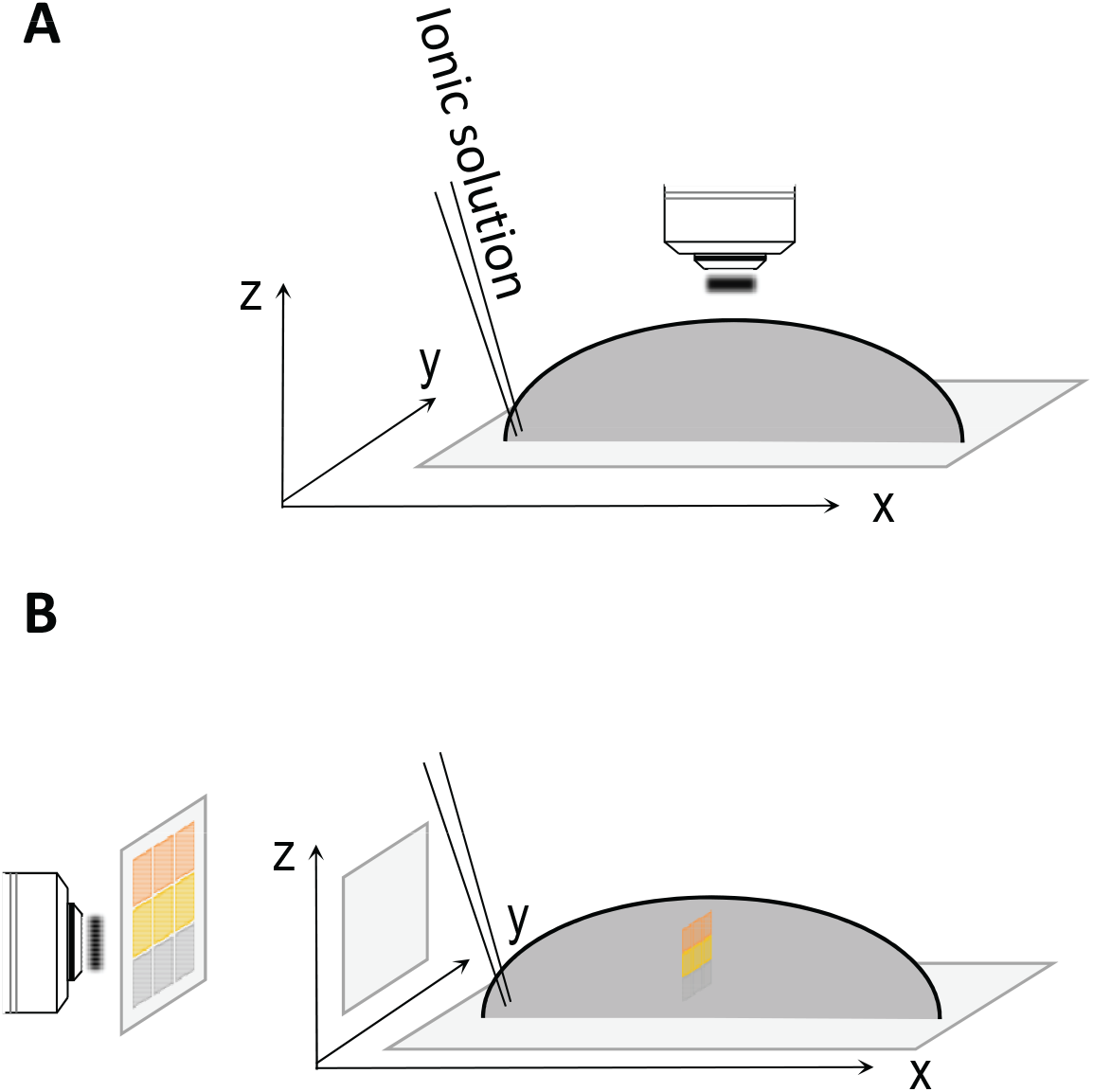
Schematic views of the droplet assay. Potassium was applied at the base of each droplet containing zoospores and at a point of the circumference. Subsequent characterization of metric of zoospore motion and potassium diffusion was performed based on micrographs generated either in the horizontal (***A***) or vertical (***B***)planes.

**Figure 2:**
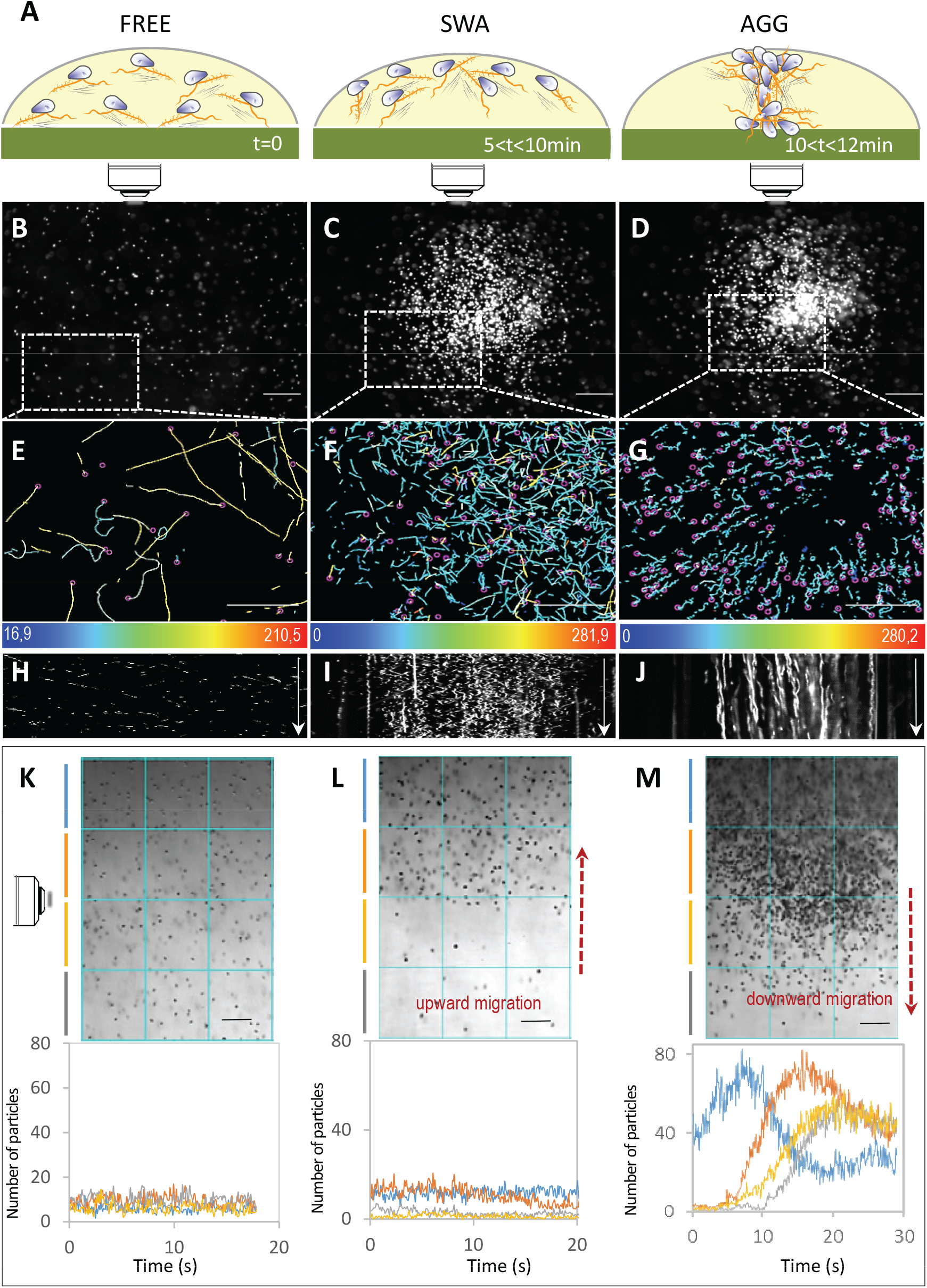
Zoospore motion in response to K^+^ application. **(A)**Summary scheme of distribution and displacement of cell population occurring during three defined different states: **FREE** for zoospores swimming freely and distributed randomly before gradient sensing; **SWA** and **AGG** respectively for zoospores forming a swarm and then an aggregate upon gradient sensing. The range is indicative of the time at which each sequence occurred after KCI application, (***B***) to (***J***) illustrate patterns observed in the horizontal plane for zoospore distribution (***B***, ***C***, ***D***), swimming paths (***E***, ***F***, ***G***) and 2D-kymographs drawn for 10 s (***H***, ***I***, ***J***). Swimming paths were recorded for xy coordinates corresponding to the areas delimited by dotted rectangle in ***B***, ***C*** and ***D*** during 2s for (***E***, ***F***) or 4s for (***G***). The color-coded stands for mean velocity information with range limiting values indicated at the bottom of each inset in μm/s. On the kymograph image, the vertical axis represents time (white arrow), the horizontal axis corresponds to the pixel’s intensities along the length of the selected line: a line ROI of **1500** μm located at the center of the field along the x axis. (***K***, ***L***, ***M***) illustrate distribution and displacement patterns observed in the vertical plane. Upper insets correspond to representative micrographs for FREE SWA and AGG. Lower insets show histograms of zoospore number per unit area, over 20 or 30s and at each depth identified by the color code in the 3 states. SWA is correlated with an upward migration (signified by ascending red arrow) while AGG is correlated to downward migration (descending red arrow). The bi-directional and sequential movement of cell population results in a 5 to 7-fold increase of local cell density during downward migration (compared ***K*** to ***M***). Values of zoospore number are the mean calculated from 3 the consecutive unit areas and based on numeration of particles with size ranging from 3 to 30 μm from converted binary images. Bar sizes: 100 μm

The process occurred with potassium and sodium, whatever the associated anion (chloride, acetate, permanganate, sulfate) and within the pH range from 5 to 8. Among other cations, only H^+^ application induced similar effects in the tested conditions although H^+^ treatment did not provoke aggregation: application of 0.001 to 0.01 μmoles triggered local but transitory gathering of zoospores; higher amounts promoted ring formation before extensive cell death.

Aggregation maximized within 12-15 minutes when at the starting point 0.5 to 3 μmoles of potassium were applied per 100 μl of cell suspension (Movie S1, Figure 2A-D). Assuming that ionic diffusion flow causes the ionic concentration to tend towards homogeneity, this led at the equilibrium point to a final range of concentrations of 5 to 30 mM. Lower amounts had no apparent effect on zoospore motion, while higher ones provoked local cell gathering but no aggregation and high rate of cell death by osmolysis (not shown). Na^+^ application induced aggregation but at a higher range than K^+^ (application range: 5 to 30 μmoles; final concentration: 50-300 mM). K^+^-induced-aggregation was observed for cell density extended from 10^5^ to 4.10^6^ zoospores/ml with a quorum ranging from 5.10^3^ to 2.10^4^ cells. For further analyses and in the following, the assays were performed at pH 6.5 by application of 0.5 μmole of KCl per 100 μl of cell suspension at 5.10^5^ cells/ml.

### 3.2. Behavior of zoospores sensing potassium

In the FREE state, both horizontal (Figure 2E) and vertical (not shown) trajectories were determined. A kymograph (Figure 2H) gives the picture of moving zoospores (appearing as oblique lines) in the horizontal plane, evolving at a velocity V_xy_ of 164 ± 44 μm/s (Table 1). Measurement of V_XZ_ indicated that velocity (129 + 39 μm/s) was also homogeneously dispersed (not shown).

**Table 1.** Mean speed of zoospores

| Velocity | FREE | SWA | AGG |
| --- | --- | --- | --- |
| $V_{xy}$ | $164 \pm 44 \mu\text{m/s}$ | $112 \pm 49 \mu\text{m/s}$ | $24 \pm 9 \mu\text{m/s}$ |
|  | Seq1 n=129 | Seq1 n=96 | Seq1 n=157 |
|  | Seq 2 n=126 | Seq2 n=102 | Seq 2n=155 |
|  | Seq3 n=137 | Seq3 n=99 | Seq3 n=93 |
| $V_{xz}$ | $129 \pm 39 \mu\text{m/s}$ | $146 \pm 42 \mu\text{m/s}$ | $38 \pm 24$ |
|  | Seq1 n= 187 | Seq1 n=222 | Seq1 n= 25 |
|  | Seq n=208 | Seq2 n=247 | Seq n=14 |
|  | Seq3 n=198 | Seq3 n=176 | Seq3 n=11 |
|  | Seq 4 n=219 | Seq 4 n=185 |  |

In the SWA state (3 to 10 min), the trajectories in the horizontal plane were undirected but limited to the geometry of the swarm (Figure 2C, F). Compared to the FREE state, V_xy_ in SWA state was reduced and more variable (112 ± 49μm/s), with a decrease especially at the swarm periphery as illustrated on the kymograph by “low”-sloping of lines at the edge of the swarm. Static peripheral zoospores appear as vertical lines parallel to the time axis, while moving ones within the swarm appear as tilted lines (Figure 1). Moreover, swarm formation was correlated with a continuous upward migration (Movie S2) that culminated with accumulation of almost all cells at the air-liquid interface, and with only very few cells detected below in the scanned area (Figure 2L).

When observed in the horizontal plane at the AGG state (Figure 2D, G, J) (t=10 to 12 min) all cells undergo migration with centripetal trajectories in the horizontal plane (Figure 2G), at very low velocity V_XY_ (24 ± 9 μm/s) towards a restricted area (1 to 5.10^6^μm^2^) at which zoospores stop their linear movement and form an aggregate (Movie S1, Figure 2D). They appear as static object on the kymograph (Figure 2J). Once plume was constituted, cells abruptly underwent downward migration leading to sequential redistribution of zoospores according to a downhill gradient along the vertical axis (Figure 2M, Movie S3). Redistribution of zoospores along the vertical axis was correlated with number of cells reaching a mean V_XZ_ of 38 ± 24μm/s in the vertical plane (Table1). This value was different of those calculated in the other situations analyzed in this study: zoospores swimming freely (128 ± 35 μm/s) or above front migration (159 ± 52 μm/s), cells floating below front migration (8 ± 13 μm/s), zoospores fixed with paraformaldehyde and only submitted to gravity and buoyancy (138 ± 19 μm/s). At the end of the sequence, zoospores were living and motile (Movie S4), exhibiting flagella (Figure S1) before progressive and massive encystment.

Obviously, displacement of plumes exhibited a high vertical component during down migration (Movie S5; Figure S2). However, at zoospore population level trajectory and velocity patterns were heterogeneous both in space and time. Velocity patterns revealed higher velocity for zoospores both in the front of migration and in more central location in the heart of the plume at the beginning of sedimentation (5s<t<20s). Then, distribution of higher velocities was limited to central location (20s<t<25s) Figure 3A). At this step, analysis of individual behavior of zoospore was difficult because of high cell density. The vertical component of trajectories was revealed only when individualization of particular zoospores could be achieved for extended time (Figure 3B). For each cell analyzed, the downward migration effectively resides at first in a virtually rectilinear and vertical displacement, including within an approximate period length of 10-12 seconds. The vertical velocity then reached 50-52 μm/s. A second component of the displacement follows, consisting of a modification of trajectory patterns which then becomes helical and a vertical velocity which is highly reduced probably when cells reached the threshold concentration range defined below (Figure 3B).

**Figure 3:**
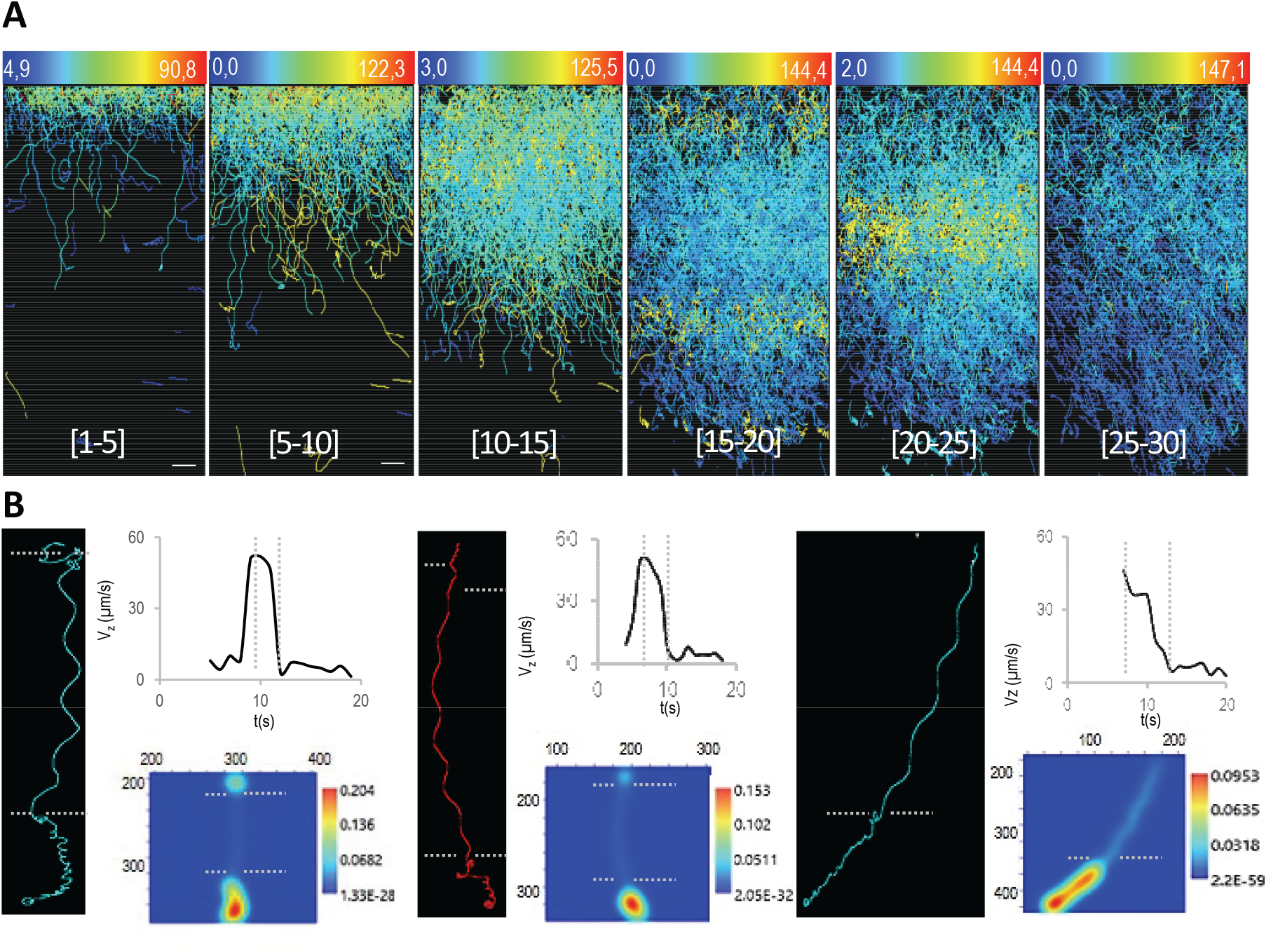
Downward migration. (***A***) Evolution of trajectory patterns during downward migration in the AGG phase. The numbers in the square brackets indicate the window time of 5 s elapsed from the onset. The color-coded stands for mean velocity information and range limiting values are indicated at the top of each inset (μm/s). (***B***) Velocity parameters for three representative zoospores. The left panels show trajectories tracked for at least 15 s. Dotted lines delineate distinct facets in particular the vertical pattern of downward migration and the linear/helical transition discernable at low part of paths. The upper right panel shows kinetics of vertical velocity. The lower one shows the map of point density generated from x/z coordinates of each of the 3 trajectories using the gaussian kernel density tool of PAST3 [29]. Scaling gives an estimate of the number of points per unit area. It should be noted that the low point density is correlated with high vertical velocity. The figure shows the data for a representative experiment. Bar sizes: 100 μm.

### 3.3 Zoospores sensing potassium are distributed according to the ionic gradient

The droplet assay led to heterogeneous and uncontrolled gradient concentration and was inappropriate to analyze relationships between zoospore and K^±^ extracellular gradient distribution. To go further in understanding how zoospores behave toward various concentrations of potassium we generated a diffusion gradient in a millifluidic device (Figure 4A) consisting basically in one channel, one inlet and one outlet. The cells were loaded into the channel (l:17 mm; w: 3.8 mm; h: 400 μm) in the presence of APG-2, a K^±^ sensing probe, prior to a spot application of KCl at the inlet. The fluorescence dynamic range of APG-2 upon ion binding and the evolution of zoospore distribution were followed along the channel by confocal microscopy (Figure 4A). The fluorescence intensity was measured for determining K^+^ concentration which is plotted in Figure 4B. The measured fluorescent concentration profile indicated that the dynamic range of K^+^ concentration that could be resolved in this device were respectively [0-17mM] at 1, 3, 5 or 10 min of diffusion. A slight minority of the cell population was unable to swim mainly at immediate vicinity of the application spot. The vast majority of zoospores was not altered in motion (velocity and random trajectory, data not shown) but had a restricted distribution. Cells were submitted to negative chemotaxis [6] and found optimum conditions for their swimming away from the higher ionic concentrations. The no-swimming zone expanded over time as the ionic diffusion progressed. The measurement of the APG-2 fluorescence intensity led to determine the range of higher concentrations consistent with the movement of zoospores at different time points (Figure 4A and C). The cell distribution depended on the local K^+^ concentration. The variation of this concentration at the frontier between the swimming and no-swimming zones covered the narrow 1–4-mM range according to the different time points. There was no significant change for this range over time. The chemotaxis assay was monitored for a period of thirty minutes without observing changes in the characteristics of zoospore motion in the appropriate swimming zone except their inability to evolve in the high potassium concentrations. This indicated that K^+^-negative chemotaxis may initiate but was not sufficient to induce aggregation in these conditions.

**Figure 4:**
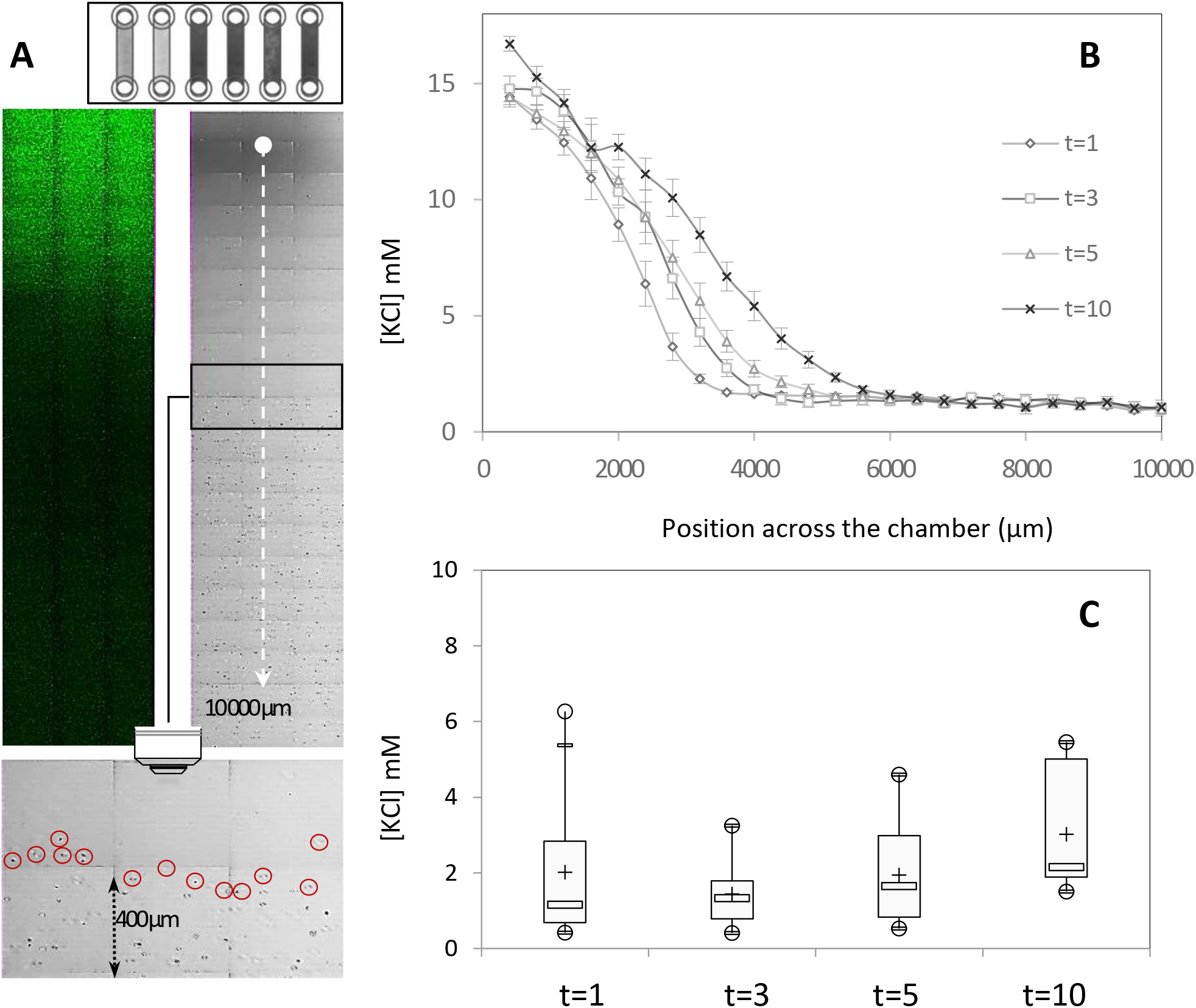
A potassium gradient drives zoospore distribution in a passive dispersion system. (***A***) Zoospores observed in bright field (right panel) and APG-2 fluorescence emission pattern captured by confocal laser microcopy (left panel) in a millifluidic device. The point blank, right between well and chamber, indicates site of application of 1 μl of 0,5M KCI at t=0; the white dotted arrow shows the overall distance screened for analyzing both zoospore distribution and APG-2-fluorescent signal at different time points (1, 3, 5 and 10 min). The lower panel illustrates the location of a 0.48 mm^2^ area used for the determination of the upper position in the chamber suitable for zoospore motion. Encircled cells indicate at time t, the location of those that schematically delimited the superior range of the gradient suitable for zoospore exploration, (***B***) Measurement of fluorescent intensity across 10 000 μm of the chamber defining the concentration profile which exhibits a non-linear gradients pattern. (***C***) Box plot of the higher ionic concentration suitable for zoospore motion. Minimum and maximum values are depicted by white dots; the box signifies the upper and lower quartiles, the mean and the median are represented respectively by ± and a white rectangle within the box for each raw data. Values are the means of 5, 6, 5 and 4 replicates for 1, 3, 5 and 10 min, respectively.

### 3.4. Changes in zoospore motion in response to K^+^ in a microfluidic device

To explore K^+^-induced behavior at the single cell level, zoospores were submitted to different gradient of KCl in a microfluidic system as explained in the method section. Zoospores were injected in the central inlet into a chemotaxis observation chamber (H x W = 0.2 mm x 1 mm), with the two side channels being perfused with 100 mM KCl (Figure 5A). The following cycles were performed: Zoospores and KCl were first injected together; at t=0, the flow of KCI was stopped during a time T (varying from 5s to 15s) to partly flush the KCl; then the flow of zoospores was also stopped, and the movement of the zoospores was observed during at least 20s. A snap shot of the chamber is shown in Figure 5B (upper picture).

**Figure 5:**
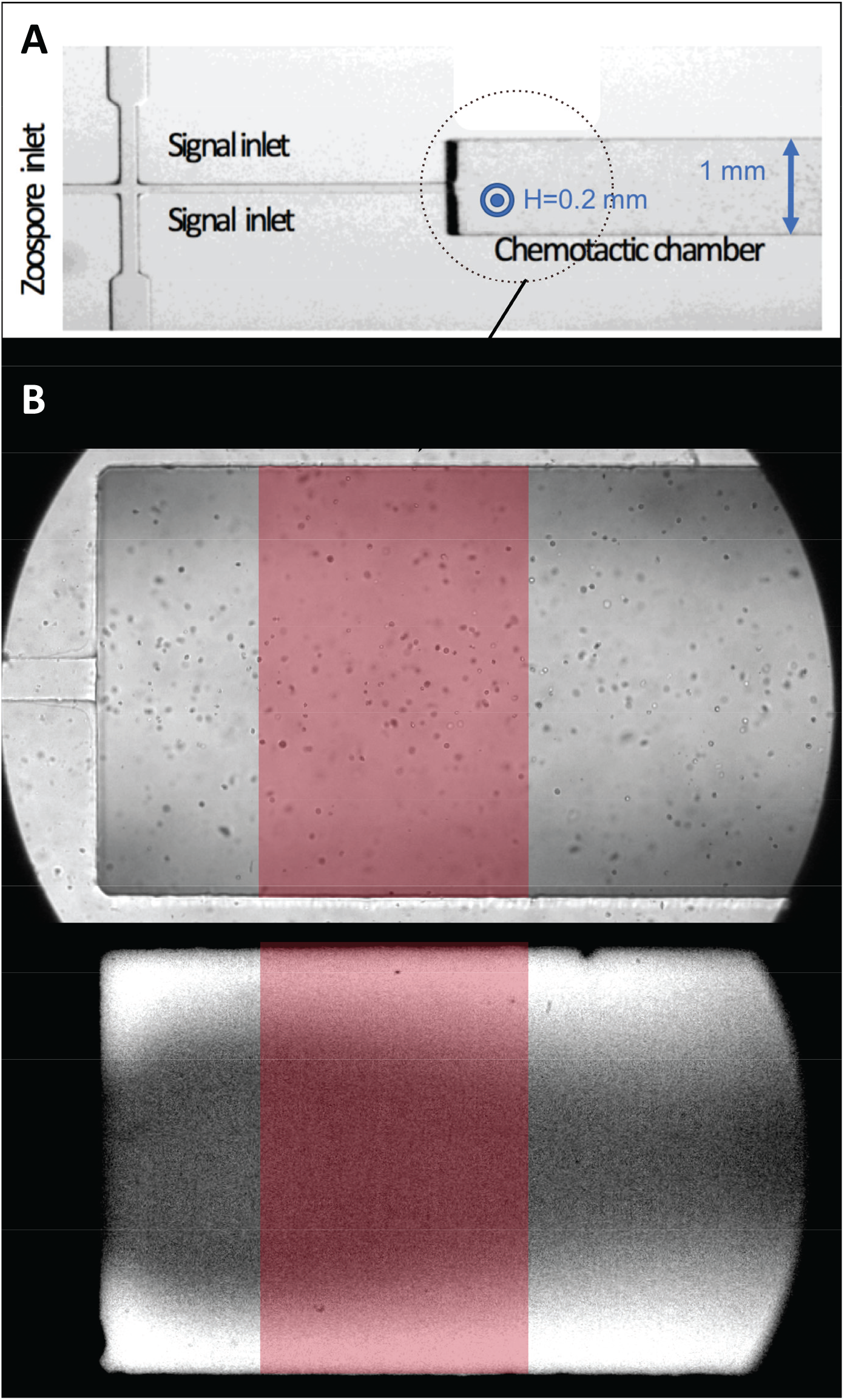
Microfluidic device. (***A***) Device picture showing the three inlets and the chemotactic chamber. (***B***) snap shots of the chamber in bright field (upper), showing zoospores, and in fluorescence (lower), allowing to map the concentration of potassium in the chamber using fluorescence of APG-2 probe.

In another experiment without zoospores, using the same set-up, a few cycles as described above were performed with KCl mixed with APG-2 to get the spatiotemporal map of the potassium concentration in the chamber. Due to the geometry of the microfluidic set-up, the KCI is concentrated in the upside and downside (on the image) regions of the chamber as shown on a snap shot in Figure 5B (lower picture).

To characterize the migratory process, we tracked cells from t=0 to t=20s to extract their trajectories, calculate the local density of cells, and measure the speed of individual cells. The area used for this analysis was restricted to the red window shown in Figure 5B. The same area was selected to calculate the spatiotemporal potassium concentration. Both density of spores and concentration of potassium were obtained by sliding window method (see §2.4.), by calculating the mean values in rectangular windows sliding along the direction of the potassium gradient (the Y coordinate in our set-up).

The results are shown in Figure 6. When the KCl was flushed during 5s, a large majority of zoospores quickly adopted circular trajectories (Figure 6A and Movie S6) at low speed, below 40 μm/s (fig. 6D and 6G). The residual concentration of potassium was thus supposed to be higher than the threshold of 3-5 mM, and to “freeze” the cells immediately. The concentration profiles in Figure 6D and 6G are in agreement with this interpretation: after 4s, potassium was above 5 mM in more than the half of the observed area, and after 16s, the concentration was above 10 mM in the whole area.

**Figure 6:**
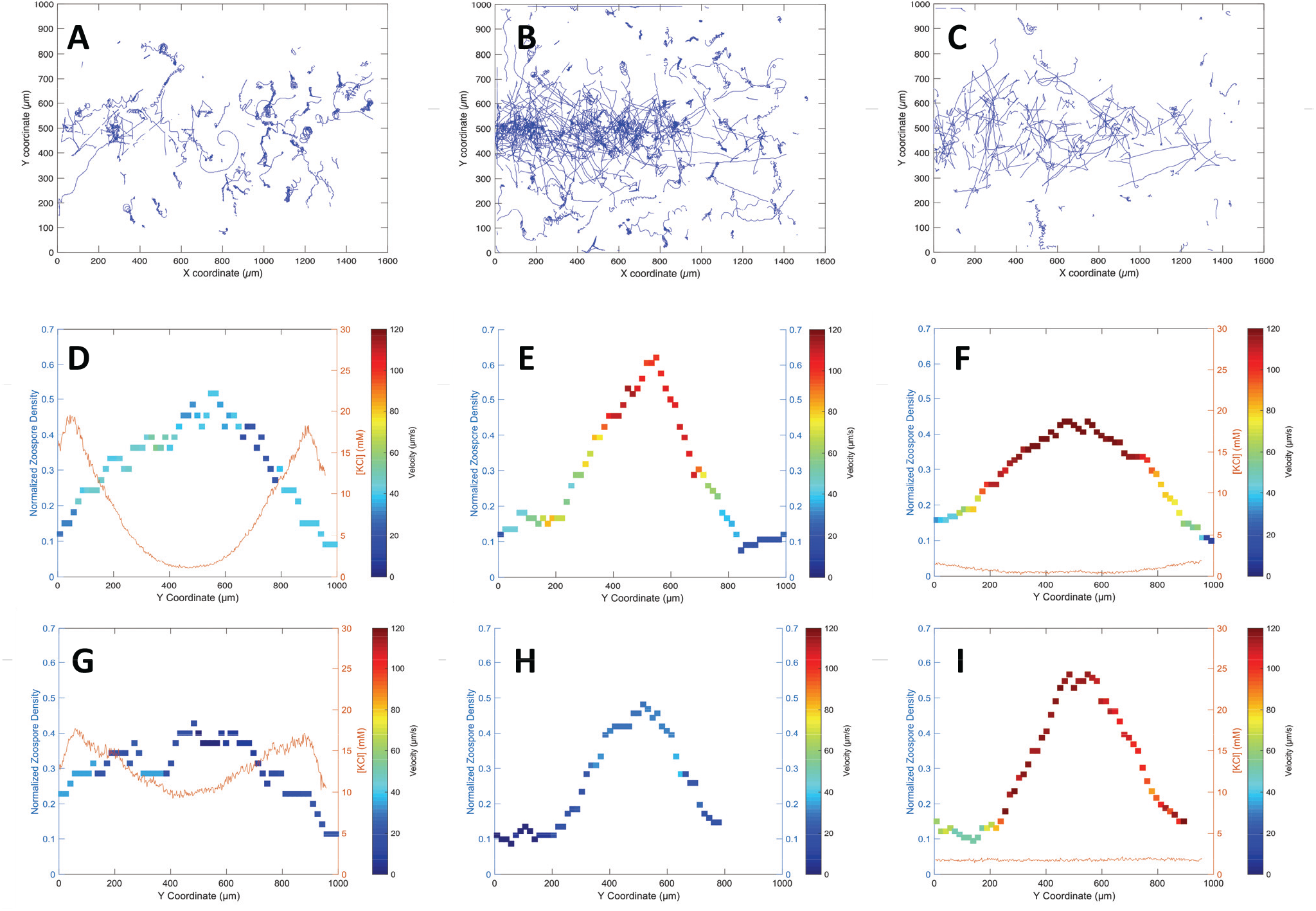
Trajectories, density profiles and speed of the zoospores in the chamber. Trajectories of the zoospores in the chamber after KCI flushing time equal to 5s (***A***), 10s (***B***) and 15s (C, only one tenth of trajectories shown for convenience). (***D-E-F***): Local density of zoospores (squares) with color map for the associated velocity, and potassium profile (red lines), after 4s. (***G-H-I***): same representations after 16s.

When the KCI was flushed during 10s, some zoospores first moved to the central zone (Figure 6B and Movie S7), as expected to escape upside and downside zones at high potassium concentration. They had linear trajectories and moved at relatively high speed (above 100 μm/s) after 4s (Figure 6E). Nevertheless, some cells initially in upside and downside zones adopted immediately circular trajectories with low speed (Figure 6B and 6E), suggesting that the concentration of potassium was too high to allow the cells to escape these zones. After a dozen of seconds, all the cells shifted to low speed-circular trajectories, suggesting that the potassium had diffused until the central zone and had reached the threshold concentration. Unfortunately, we were not able to measure the potassium concentration during this cycle. After 16s, in the central zone, the speed of the cells was below 40 μm/s (Figure 6H), showing that they had shifted to low speed-circular trajectories.

When the KCI was flushed during t=15s, the behavior was close to the observations made for T=10s, except that the area where the zoospores moved with linear trajectories at high speed was greater (Movie S8, Figure 6C and 6F), and that after 16s, the cells went on to move with high speed-linear trajectories (Figure 6I). This suggests that the concentration of potassium was sufficiently low to allow the cells to move. The concentration profiles after 4s and 16s (Figure GF and 6I) are in agreement, as the concentration is always below 3 mM. Interestingly, it can be noticed that: (i) between, 4s and 16s, the cells tend to migrate to the central zone, although the potassium is at very low concentration; (ii) after 16s, the concentration profile seems to be flat, whereas the density of zoospores is peaked. These observations suggest that a motion leading to aggregation could be trigger here by a cue different from potassium concentration. More investigations need to be performed to shed light on this behavior.

## 4. Discusssion

It has been previously shown that *Phytophthora* auto-aggregation results from bioconvection pattern formation [17] or from bioconvection combined to positive chemotaxis [13]. It has also been demonstrated that *Phytophthora* potassium sensory cues regulates zoospore behavior [5, 6]. In this report, we present *in vitro* evidence that potassium gradient sensing by *P. parasitica* zoospores is a primary stimulus, which induces a synchronized zoospore behavior and cell aggregation. The macroscopic pattern is reminiscent of the auto-aggregation one. The cell behavior evolves during a single bi-phasic temporal sequence different that previously described for auto-aggregation.

### 4.1. Negative chemotaxis and bioconvection

At first, negative chemotaxis to potassium leads the cells to go to region where potassium concentration is below the threshold range of 1-4 mM. Then, the spatial concentration of cells arises as a result of the geometric arrangement. In a droplet, negative chemotaxis enforces cell concentration through upward migration toward upper surface of the suspensions. Once the zoospore density exceeds a certain threshold bioconvection occurs. The suspension surface becomes too dense and zoospores constitute plumes which undergo downward migration. No apparent positive chemotaxis between formed plumes [13] was noted. However, the distance separating plumes from each other was variable (≈0,5 to 3mm) rendering difficult the quantification of this parameter in the tested the conditions. At cellular level, each of the two sequences is characterized by distinct motion parameters (upward cell concentration and plume downward migration). The congruent results emerging from the droplet (in vertical and horizontal planes) and microfluidic (horizontal plane) assays lead to the following scheme. As long as K^+^-treated cells have the opportunity to escape from high concentration areas, they exhibit linear trajectory and turn back toward the areas below the threshold concentration of 1-4mM, this without measurable interference on velocity (Figure 1, 2, 4). When the conditions force the cells to be located beyond this threshold (Figure 3), two drastic changes are observed: an important reduction of velocity and a switch from a linear trajectory to a rotational one. Thus, in most cases, the K^+^-induced aggregation phenomenon is observed when the motion of a quorum of zoospores (5.10^3^-2.10^4^) explores territories which are delimited by the concentration field and its spatial distribution in the different devices used here. Although, in the microfluidic devices, it seems that zoospores can stay longer in an aggregate states when spatiotemporal variation ok K^+^ gradient are higher, leading to the hypothesis of a higher response mediated by a signaling behavior between zoospores enhancing their response.

### 4.2 Sources for K^+^ gradient generation in natural environment

The *in vitro* induction by K^+^ of zoospore gathering demonstrates that aggregation may result from perception of an external signal(s). How such gradients can be achieved in natural habitats? At first glance, they may be generated by efflux mechanisms from zoospores, release from microbiota sharing the same biotope, rhizospheric activity and/or exchange dynamics in soil.

Self-generation of K^+^ by a net K^+^ zoospore efflux seems highly unlikely. In freshwater, the osmolality of the zoospore cytosol is always higher than that of the external environment. Throughout the course of their displacement, a major challenge of (wall-less) zoospores is the removal of excess of cytosolic water rather than ions for maintaining homeostasis. The water excess is collected into the contractile vacuole complex (CVC), the osmoregulatory organelle, and discharged to the extracellular environment [9]. Calculating the K^+^ diffusion potential from cells also rejects the K^+^-zoospore-efflux hypothesis. If we consider high densities (10^6^-10^7^/ml) of zoospores (diameter of 10 μm), and K^+^ intracellular concentration ranges of 100-200 mM, then the relative volume expansion rate of the cell population is fluctuated from 0,052 to 0,52%. This would lead to a K^+^ extracellular concentration between 0,05 and 1 mM in the incongruous case that the whole cell content of potassium would be released by efflux. Thus, it is not plausible to reach by this way the concentration threshold of 1-4 mM determined in this study as a stimulus eliciting a pattern reminiscent to auto-aggregation. As in the case of most prokaryotic and eukaryotic cells [24], we can rather speculate that depolarization due to the increase in extracellular potassium concentration leads to fluctuations in the membrane potential of zoospores that could lead to modifications of flagellar beating, cellular responses such as osmoregulation and/or cell-to-cell signaling.

To what extent K^+^ release by microbiota sharing the zoospore habitat could mediate their aggregation remains to be evaluated. Potassium appears to be key in the displacement of bacteria, in the physical composition of microbiota [24] and in the process of pathogenesis [25]. Electrical signaling mediated by potassium ion channels regulates cell-cell dialogs within a bacterial biofilm with potassium driving attraction of distant cells of different species [24]. The range of bacteria-generated gradients that are effective for attracting prokaryotes *in vitro*, is of the same order of magnitude as that impacting the movement of zoospores in our experimental scheme. Such gradients could therefore have an impact on the distribution of *P. parasitica* propagules through repulsion.

Regarding rhizospheric activity and soil dynamics exchanges, obviously any area that tends to generate elevated K^+^ gradient will also tend to be repelling for *P. parasitica* zoospores. In soils, the total K^+^ contents generally range between 0.4 and 30 g kg”^-1^ [26] with two components. The soil particles, which bond about 98% of total K^+^ content, can constitute main obstacles for zoospore tracking. The surrounding film of water (2% of total K^+^ content) exhibits an approximate potassium concentration range of 0.2-15mM, that reaches 5-10mM in most fertigated agricultural soils [26]. Thus, it is reasonable to speculate that K^+^ exchange dynamics in soil constitutes an important parameter for zoospores distribution and aggregation. At the scale of an infected host plant, zoospores repelled by soil particles and constrained in their movement in the film of water should be prone to move towards host tissues. This inclination should be reinforced by the root K^+^ uptake causing the formation of a depletion zone around the root surface [27], and by root exudates which attract zoospores [1, 28]. Thus, the physical and chemical distribution of K^+^ at the soil-root interface is a parameter propitious to constitution of high density inoculum or biofilm formation on the plant surface. However, its contribution remains to be established compared to other ion dynamic exchanges such as soil acidification by roots. A more holistic view of the relationships between the concentration of various ions and zoospore distribution is required to delimitate the influence of the K^+^ one.

### 4.3. Concluding remarks

In this study, we show that potassium gradient sensing induces a synchronized zoospore behavior (due to negative chemotaxis) and cell aggregation of *P. parasitica* zoospores. In all various *in vitro* experimental setups that have been used (droplets assays, milli- and microfluidic devices), zoospores present such common behaviors, showing a clear, common response to Potassium. The use of these different setups also enlightens particularities linked to size and geometrical effects of each device. Bioconvections patterns that follow aggregation are only observed in droplet assays. In millifluidic devices, clear and quantitative response to Potassium gradient is evidenced. The microfluidic devices allowed faster spatiotemporal variations of Potassium concentration which allow to observe an aggregation process which seems to constitute a stronger response compare to the one expected for quasi-static negative chemotaxis. In other words, even when the Potassium concentration drops down to small values, that zoospores should explore, they stay in an aggregated state. Hence, perspectives of this work in that direction will consist in looking for signals sent by zoospores in presence of K^+^ that would increase their response in term of auto-aggregation behaviors. More complex microfluidics devices together with a modeling approach are being developed with this aim and will be the topic of future studies.

## Supplementary data legends

**Figure S1:**
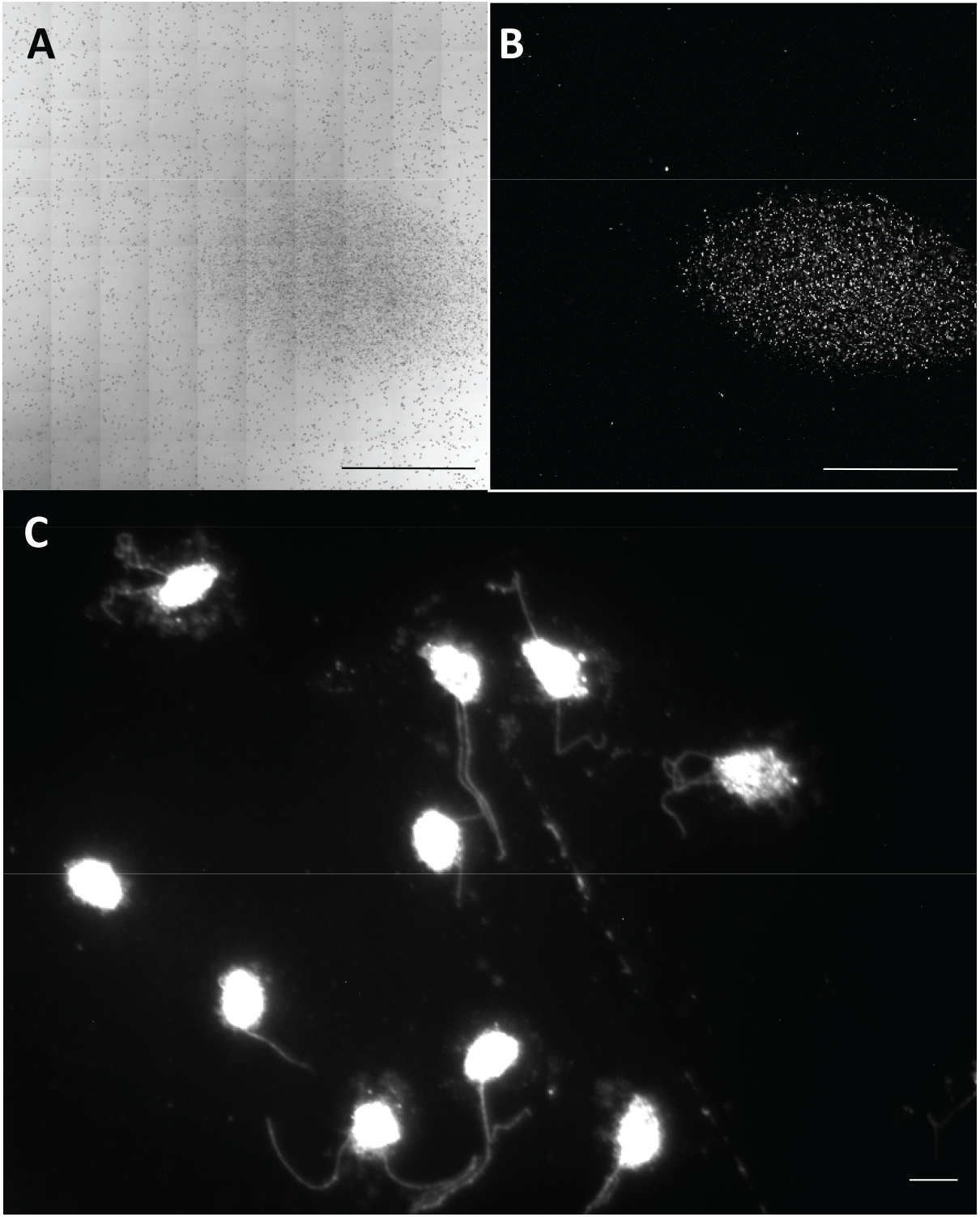
Distribution of K^+^-treated and aggregated zoospores. Before treatment with potassium, zoospores were stained with 0,001% Nile Red and mixed with polystyrene 10μm microspheres. After treatment (t=20 min), the microspheres+zoospores distribution were observed with a bright-field transmitted light detector (***A***). The zoospores stained with the fluorescent dye were observed with 514 nm excitation, ranging from 534 to 700 nm (***B***). The comparison between ***A*** and ***B*** illustrates the homogenous repartition of microspheres while aggregated zoospores are restricted to an area of 1,8 10^6^ μm^2^. (***C***) A micrograph showing biflagellate zoospores fixed in 1% paraformaldehyde immediately after aggregation and then stained with 0,001% Nile Red. Bar sizes 1000 μm in ***A*** and ***B*** 10 in μm ***C***

**Figure S2:**
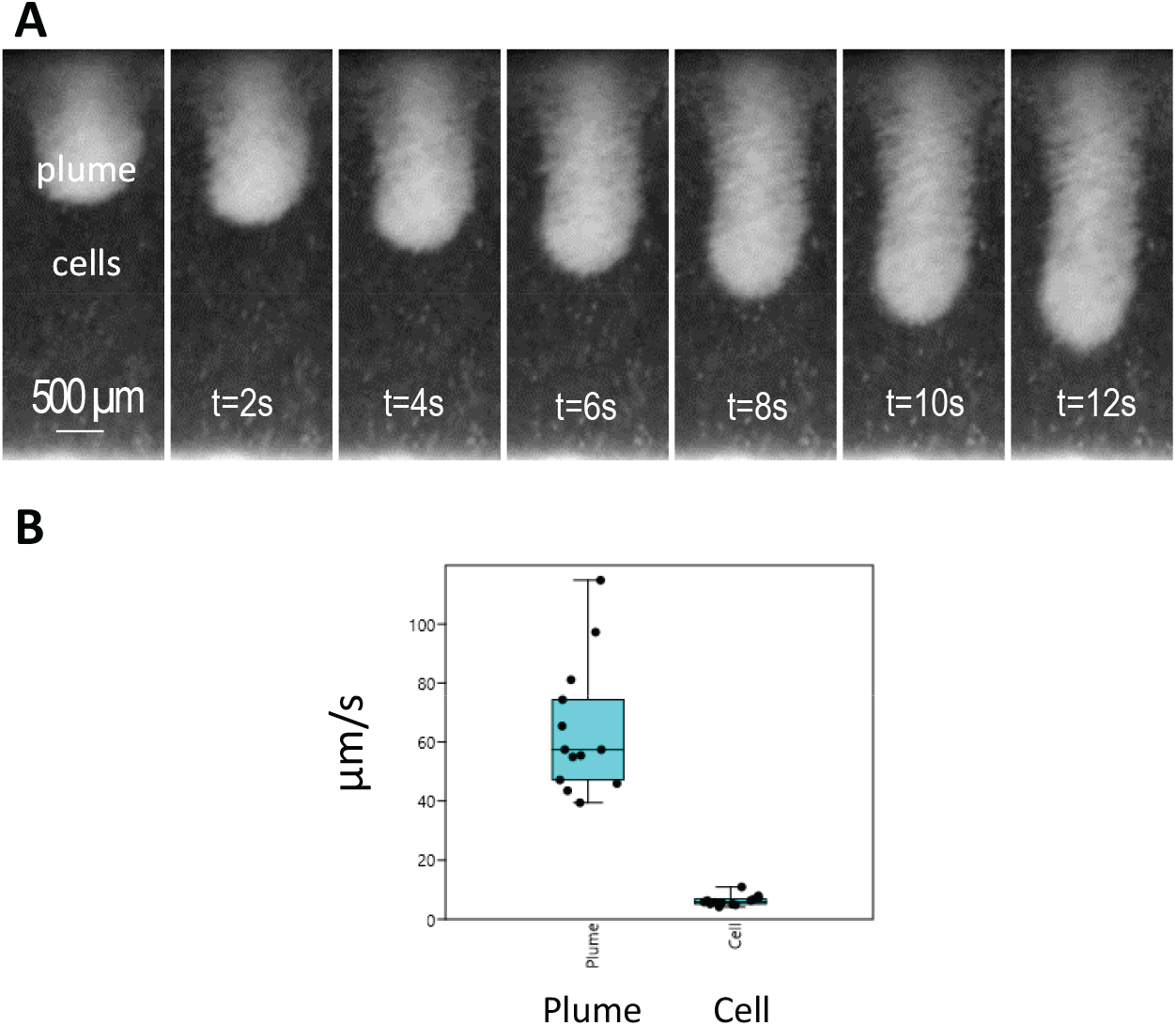
Plume sedimentation. (***A***) Time series of selected views illustrating the vertical evolution of a plume overtime, (***B***) Box plot of the sedimentation rate measured for plumes (at sedimentation front) and floating isolated cells.

**Movie S1:** Sequence of events leading to zoospore aggregation observed by dark field and upright microscopy (microscope Zeiss Z1, objective 2.5x).

**Movie S2** Upward migration observed with a vertical mounted microscope (AxioSkop Zeiss, Objective 4x)

**Movie S3** Downward migration observed with the vertical mounted microscope

**Movie S4** Cell population observed 30 min post K^+^-treatment with both motionless cysts and zoospores mainly exhibiting an anticlockwise rotation movement.

**Movie S5** Vertical displacement of plumes during down migration

**Movie S6** Zoospore behavior following KCl flushing during 5s

**Movie S7** Zoospore behavior after KCl flushing during 10s

**Movie S8** Zoospore behavior after KCl flushing during 15s

Authors’ contributions
EG and XN designed experiments.
EG and CE carried droplet and millifluidic analyses.
XN, CC and PT carried out microfluidic analyses.
EG, XN, CC, PT wrote the manuscript.

## Acknowledgements

The authors thank the Microscopy Platform-Sophia Agrobiotech Institut-INRA 1355-UNS-CNRS 7254-INRA PACA Sophia Antipolis for access to instruments and technical advice. The authors would like to thank Fernando Peruani, Emiliano Perez Ipiña (UAD, Nice) and Laurent Counillon (LP2M, Nice) for fruitful discussions.

## Notes

**Funding:** This work has been supported by the French government, through the UCAJEDI Investments in the Future project managed by the National Research Agency (ANR) with the reference number ANR-15-IDEX-01; through the “Credits Scientifiques Incitatifs” of the University of Nice Sophia-Antipolis and the “Action Recherche” of the INRA Plant Health and Environment Division.

